# zDHHC5 expression is increased in cardiac hypertrophy and reduced in heart failure but this does not correlate with changes in substrate palmitoylation

**DOI:** 10.1101/2022.08.19.504400

**Authors:** Alice Main, Andri Bogusalvskyii, Jacqueline Howie, Chien-wen Kuo, Aileen Rankin, Francis L. Burton, Godfrey L. Smith, Roger Hajjar, George S. Baillie, Kenneth S. Campbell, Michael J. Shattock, William Fuller

## Abstract

S-palmitoylation is an essential lipid modification catalysed by zDHHC-palmitoyl acyltransferases that regulates the localisation and activity of substrates in every class of protein and tissue investigated to date. In the heart, S-palmitoylation regulates sodium-calcium exchanger (NCX1) inactivation, phospholemman (PLM) inhibition of the Na^+^/K^+^ ATPase, Nav1.5 influence on membrane excitability and membrane localisation of heterotrimeric G-proteins. The cell surface localised enzyme zDHHC5 palmitoylates NCX1 and PLM and is implicated in injury during anoxia/reperfusion. Information is lacking about how palmitoylation remodels in cardiac diseases. We investigated expression of zDHHC5 in animal models of left ventricular hypertrophy (LVH) and heart failure (HF), along with HF tissue from humans. zDHHC5 expression was rapidly elevated during onset of LVH, whilst HF was associated with decreased zDHHC5 expression. Paradoxically, palmitoylation of the zDHHC5 substrate NCX1 was significantly reduced in LVH but increased in human HF. Overexpression of zDHHC5 in rabbit ventricular cardiomyocytes was not sufficient to drive changes in palmitoylation of zDHHC5 substrates or overall cardiomyocyte contractility, suggesting changes in zDHHC5 expression in disease may not be a primary driver of pathology. zDHHC5 itself is regulated by post-translational modifications, including palmitoylation in its Cterminal tail, and we found the palmitoylation of zDHHC5 may be increased in heart failure in the same manner as NCX1, suggesting additional regulatory mechanisms such as acyl-CoA availability may be involved. Importantly, this study provides the first evidence that palmitoylation of cardiac substrates is altered in the setting of HF, and that expression of zDHHC5 is dysregulated in both hypertrophy and HF.

## Introduction

Heart failure (HF) represents a major economic and global burden affecting over 26 million people worldwide, with ∼3.6 million patients newly diagnosed every year (Ambrosy *et al*., 2014; Simmonds *et al*., 2020). All clinical therapies currently aim to reduce myocardial demand by either reducing peripheral resistance and pre-load (i.e. angiotensin-converting enzyme inhibitors (ACEi)) or ventricular after-load and remodelling (i.e. β-blockers; Shah *et al*., 2017). This largely helps to compensate for a reduction in ejection fraction and systolic dysfunction, as seen in HF with reduced ejection fraction (HFrEF) and therefore is more effective here than for HF with preserved ejection fraction (HFpEF). However, even though improved therapeutic recommendations for HFrEF significantly reduce morbidity and mortality rates in clinical trials, prognosis is still poor in this group with a trial of 40,000 hospitalised HF patients demonstrating a 5-year mortality rate of 75%, independent of LVEF (McMurray *et al*., 2014; Shah *et al*., 2017). As such, there remains a pressing and largely unmet clinical need for HF therapies.

Understanding the molecular basis of HF, in both humans and animal models, is essential to developing and directing appropriate therapies. However, this task is complicated by the fact that compensatory cardiac hypertrophy, leading to normal or even enhanced systolic function and left ventricular (LV) enlargement, often precedes decompensated HF and eventual LV dilatation and dysfunction (Kemp & Conte, 2012). The biochemical and molecular changes in both these phases of the disease are distinct, and currently available animal models are often better at modelling hypertrophy than the complex clinical syndrome of HF (Houser *et al*., 2012). Interestingly, a key component of cardiac dysfunction associated with both cardiac hypertrophy and HF is the dysregulation in expression, modification and function of proteins involved in excitation contraction coupling (ECC). Many of the pathological changes in protein function in ECC during cardiac disease can be attributed to changes in post-translational modification regulation, including phosphorylation (Ramila *et al*., 2015; Sequeira & Maack, 2018) and redox modifications like S-glutathionylation (Pastore & Piemonte, 2013). The majority of proteins involved in ECC have been reported to be regulated by the lipid posttranslational modification of palmitoylation (see review by Essandoh *et al*., 2020) with recent proteomic analysis suggesting over 450 cardiac proteins are palmitoylated (Miles *et al*., 2021). Palmitoylation is a reversible modification involving the attachment of a fatty acid (most commonly palmitoyl derived from palmitoyl-CoA) to the thiol of a cysteine residue. This has a wide range of physiological effects, including regulating the location of the palmitoylated substrate, altering substrate activity by making an area more/less accessible, or regulating protein-protein interactions. The modification is catalysed by a group of chemically related zDHHC-palmitoyl acyltransferases (zDHHC-PATs) which are located throughout the secretory pathway (Main & Fuller, 2021).

Although important in regulating ECC substrate activity, the role of palmitoylation in cardiac disease and HF remains largely unknown. There have been limited studies conducted in zDHHC knock-out animals, but zDHHCs show redundancy and therefore determining individual enzyme characteristics via this route is challenging (Main & Fuller, 2021). The most well characterised cardiac DHHC is cell surface localised zDHHC5 (see review by Woodley and Collins, 2021). NCX1 and Na^+^/K^+^ ATPase both interact with zDHHC5 after its fourth transmembrane domain, in a site that also interacts with its accessory proteins Golga7 and Golga7b (Ko *et al*., 2019; Plain *et al*., 2020; Woodley & Collins, 2021). zDHHC5 directly palmitoylates ECC substrates NCX1 and PLM (Howie *et al*., 2014; Gök *et al*., 2020). ZDHHC5 has attracted attention in cardiac physiology as it is a key contributor to massive endocytosis (MEND) of the cellular membrane in anoxia/reperfusion (A/R) injury, with DHHC5 knock-out hearts showing enhanced functional recovery from MEND (Hilgemann *et al*., 2013; Lin *et al*., 2013). In the present study, we evaluated changes in zDHHC5 expression in animal models of left ventricular hypertrophy and heart failure, as well as in human ischaemic heart failure samples. For the first time, we provide the evidence that palmitoylation of cardiac substrates is altered in the setting of HF, and that expression of zDHHC5 is dysregulated in both hypertrophy and HF.

## Results

### Remodelling of the cellular palmitoylation machinery in HF

There remains a pressing need to understand the gene and protein expression changes in the failing myocardium in HFpEF and HFrEF in order to tailor existing, and develop new, therapeutic options. Despite the knowledge that zDHHC5 is involved in A/R and regulates several important cardiac substrates, the expression or activity of zDHHC5, or any other palmitoylating or depalmitoylating enzymes, has not been investigated in a HF setting. Recently, Hahn and Knotsdottier *et al*. completed a comprehensive study of RNA transcript changes in HFpEF and HFrEF compared to organ donor controls. To investigate the relevance of palmitoylation as a modification in heart failure, using the available data, we plotted the change in relative abundance in HFpEF and HFrEF compared to organ donor controls of all available palmitoylating enzymes (zDHHC-PATs), depalmitoylating enzymes (LYPLAs, PPTs and ABHDs), accessory proteins (Selenoprotein-K (Fredericks *et al*., 2017) and Golga7 (Ko *et al*., 2019)) and proteins involved in fatty acyl CoA production (acyl-CoA synthases (ACSL) and fatty acid synthase (FASN)). Several of these were found to be both up and down regulated in the failing myocardium with some changes unique to one phenotype, including zDHHC5 which was significantly reduced in HFpEF but not HFrEF. Fatty acid and fatty acyl CoA availability have recently emerged as regulators of protein palmitoylation (Main & Fuller, 2021). For example, acyl-CoA synthase (ACSL) isoforms physically associate with zDHHC5, and acyl-CoA synthase activity is required for insulin-induced palmitoylation of NCX1 (Gök *et al*., 2022; Plain *et al*., 2020). ACSLs 1 and 4 were significantly downregulated in both HFpEF and HFrEF, with significant changes in all other isoforms and fatty acid synthase in HFpEF only (Figure 1; Hahn *et al*., 2021).

**Figure 1:**
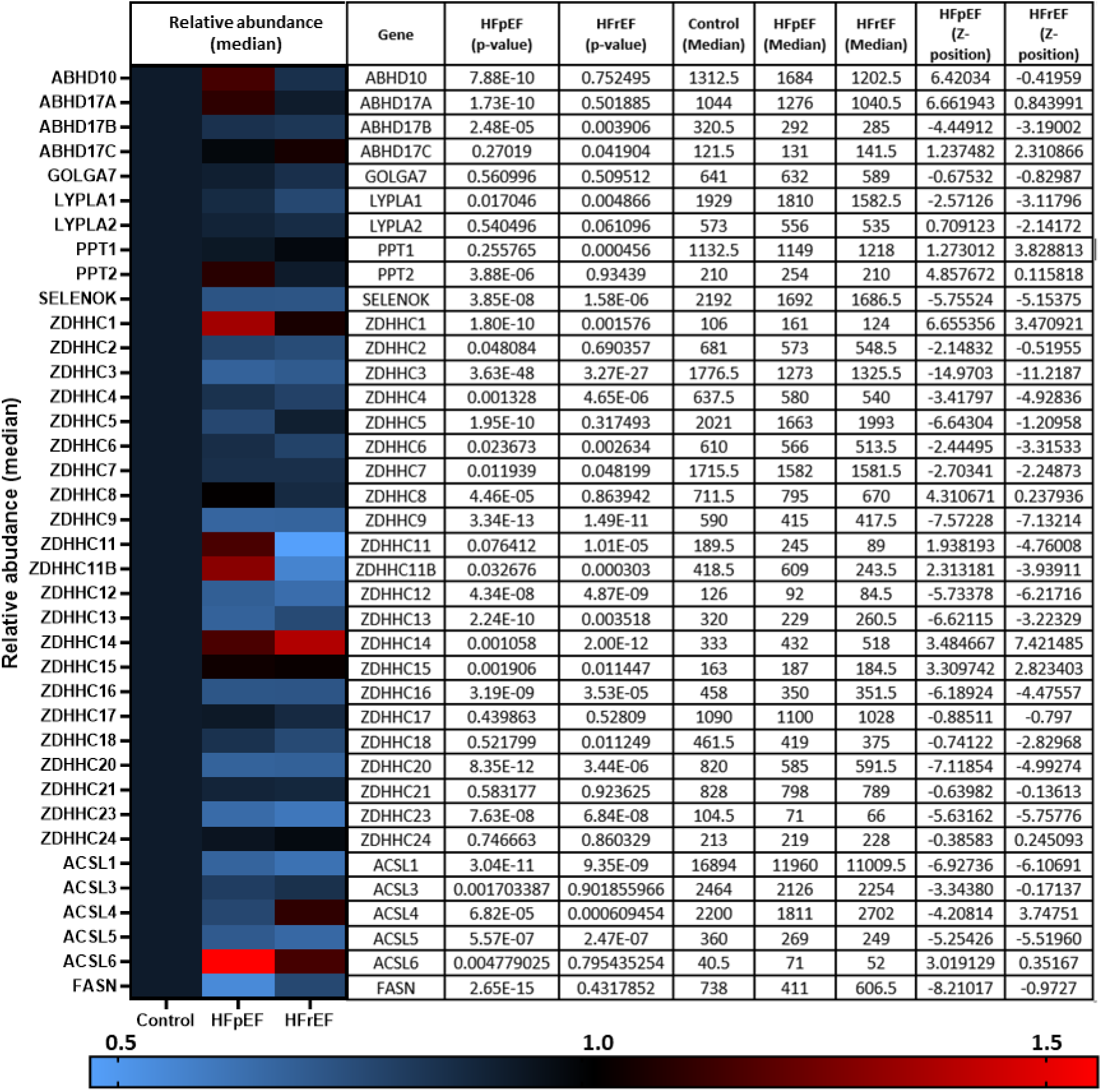
RNA transcript changes of palmitoylating and depalmitoylating enzymes and accessory proteins in heart failure. Hahn and Knutsdottir, *et al*. performed RNA sequencing on biopsies from heart failure patients with preserved ejection fraction (HFpEF, n=41), reduced ejection fraction (HFrEF, n=30) and donor controls (n=24). Abundance of transcripts of palmitoylating and depalmitoylating enzymes, including 23 zDHHC-palmitoyl acyltransferases, as well as accessory proteins and proteins involved in fatty acid synthesis, were investigated in the available data. HFpEF and HFrEF are plotted relative to control (normalised to 1) in the heat map with relative abundance and p-values detailed in the accompanying table (produced using the data repository from Hahn and Knutsdottir, *et al*. 2020).

### zDHHC5 in cardiac hypertrophy

Cardiac hypertrophy and associated remodelling of the left ventricle precedes many forms of cardiac disease, including heart failure, and is recognised as a crucial step in its pathophysiology (Rame & Dries, 2007). Left ventricular hypertrophy (LVH) induced by pressure overload via aortic constriction (‘banding’) is commonly used to investigate early molecular changes associated with pathological remodelling. This mouse model (detailed in Boguslavskyi *et al*., 2014) is associated with a 50% increase in LV mass 2 weeks after banding the thoracic aorta. We investigated changes in zDHHC5 expression associated with the onset of LVH. Compared to sham operated control mice, zDHHC5 expression was modestly increased from the earliest timepoint investigated (3 days post-banding) and was significantly elevated 2-weeks and 8-weeks after surgery (Figure 2).

**Figure 2:**
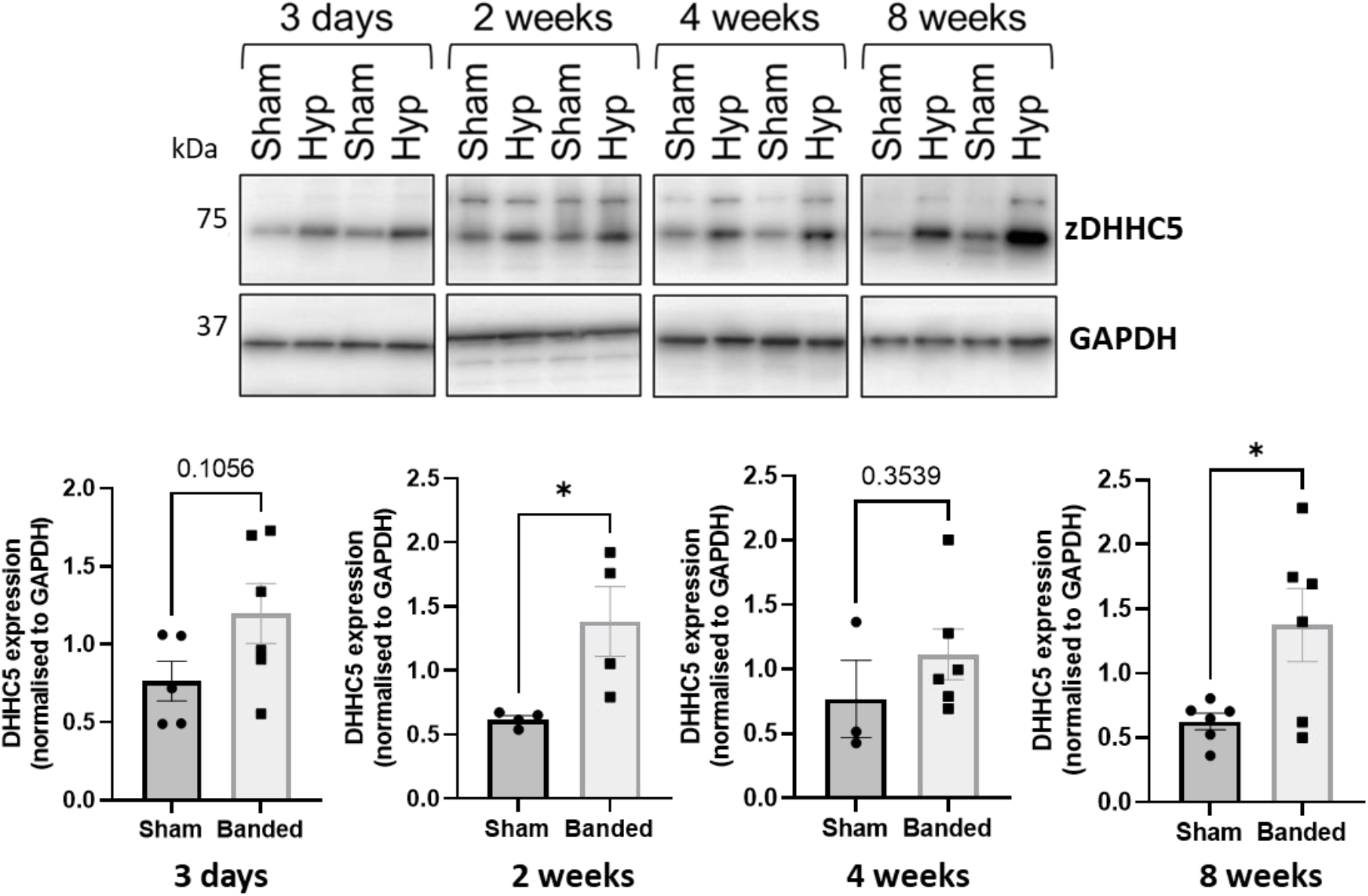
Expression of zDHHC5 is increased in mice with left ventricular hypertrophy from 3-days post-onset. Ventricular samples from mice that underwent left ventricular hypertrophy induced by pressure overload and sham controls were taken at 3 days (n=4), 2 weeks (n=5-6), 4 week (n=3-6) and 8 week (n=6) post-onset. Expression of zDHHC5 was significantly increased in hypertrophy samples compared to control at 3 days, 2 weeks and 8 weeks post-onset (*p<0.05). zDHHC5 expression is normalised to loading control GAPDH. Statistical comparisons made by unpaired Student’s t-test. Data are mean±S.E.M.

As elevated zDHHC5 expression was an early event in the onset of LVH, we investigated whether increased zDHHC5 expression is directly associated with the contractile dysfunction observed in this phenotype. We engineered adenoviruses expressing HA-tagged zDHHC5 and catalytically inactive zDHHS5, and infected adult rabbit ventricular cardiomyocytes, achieving dose dependent increases in zDHHC5 expression levels (Figure 3A). Confocal microscopy revealed localisation of HA-tagged zDHHC5 in intercalated discs, cell surface and perinuclear membrane (Supplementary Figure 1). Changes in contractile function were investigated using a CellOPTIQ® contractility system (Clyde Biosciences Ltd.), and contractility parameters measured by determining the spatial frequency of the intensity profile of sarcomere bands over time (Supplementary Figure 2; Rocchetti *et al*., 2014). Overexpression of zDHHC5 or zDHHS5 had no effect on contractile force (as determined by sarcomere shortening (Figure 3B)) or on any other parameters of contractility (Supplementary Figure 3).

**Figure 3:**
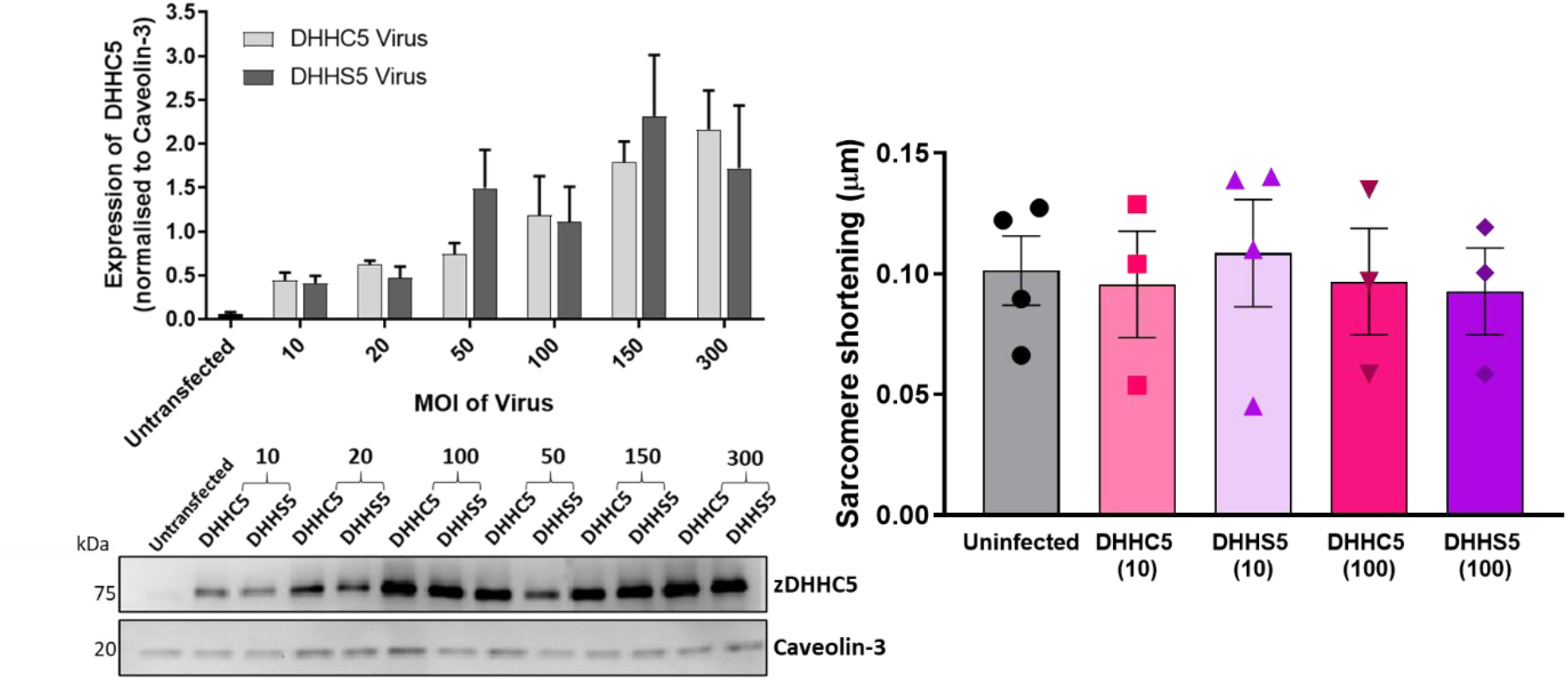
DHHC5 overexpression does not alter parameters of rabbit ventricular cardiomyocyte contractility. **A)** Rabbit ventricular cardiomyocytes were cultured for 18-24 hours in the presence of either the HA-DHHC5 virus or a HA-DHHS5 dominant-negative virus, at an increasing range of virus particles per cell or multiplicity of infection (MOI; 10, 20, 50, 150, 300). DHHC5 protein expression was determined via western blot, normalising to housekeeper protein Caveolin-3. HA-DHHC5 and HA-DHHS5 viral infection led to a dose-dependent increase in DHHC5 expression. Data are expressed as mean±S.E.M and is a representative image of n=3-4 biological replicates. **B)** Contractility recordings from cells infected with MOI of 10 and 100 were taken using CellOPTIQ®. Viral infection with either DHHC5 or DHHS5 at MOI 10 or 100 had no significant effect on altering the parameters of contractility including sarcomere shortening. Data are mean±S.E.M analysed using a one-way ANOVA with Sidak’s post-hoc test. N=3 biological replicates for S10, C100, S100 and n=4 biological replicates for uninfected and C100, with n=20-51 cells per replicate.

As both LVH and viral infection were associated with increased zDHHC5 expression, we hypothesised palmitoylation of zDHHC5 substrates NCX1 and PLM would be increased in these samples. Despite overexpression of zDHHC5, development of LVH was associated with unchanged PLM palmitoylation, whilst NCX1 palmitoylation was significantly reduced (Figure 4A). In rabbit cardiomyocytes overexpressing HA-zDHHC5, there was no significant change in palmitoylation of NCX1 or PLM compared to uninfected control suggesting increasing zDHHC5 expression is not sufficient to drive changes in substrate palmitoylation (Figure 4B).

**Figure 4:**
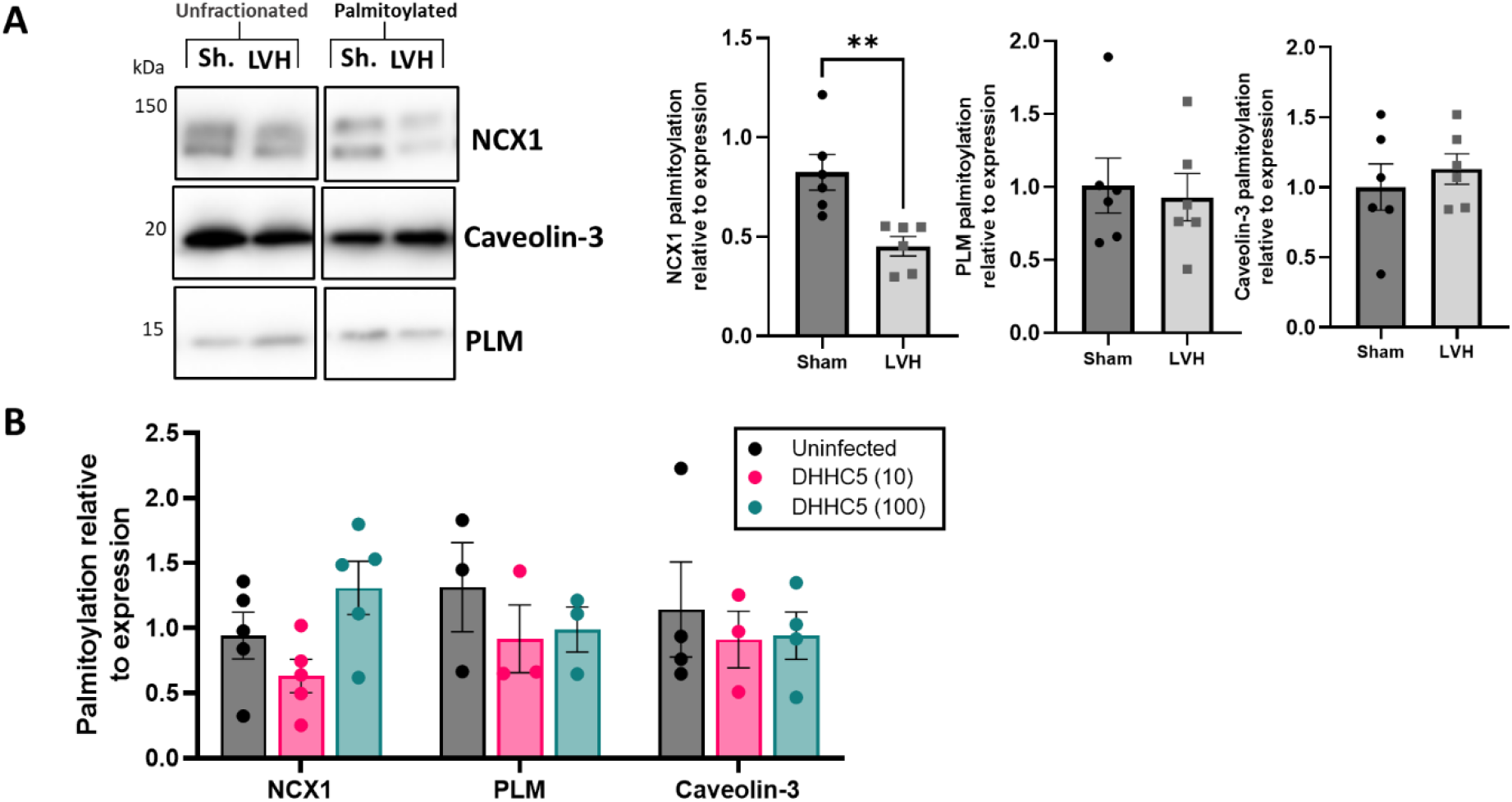
zDHHC5 overexpression in LVH or viral overexpression in rabbit cardiomyocytes does not correlate with an increase in palmitoylation of zDHHC5 substrates. **A)** Ventricular samples from mice that underwent left ventricular hypertrophy (LVH) induced by pressure overload and sham (Sh.) controls were taken at 8 weeks post-onset. Acyl-RAC revealed palmitoylation of zDHHC5 substrate NCX1 was significantly reduced and palmitoylation of PLM remained unchanged. **B)** Rabbit ventricular cardiomyocytes overexpressing HA-zDHHC5 (MOI 10 and 100) showed no significant change in NCX1 or PLM palmitoylation. Data are mean ±S.E.M analysed via an unpaired t-test (LVH) or a one-way ANOVA with a Dunnett’s post-hoc test (overexpression). *p<0.05.

### zDHHC5 in heart failure

As zDHHC5 expression is increased in early cardiac hypertrophy, we investigated whether this difference persists in HF by analysing two experimental models (rabbit and pig) of myocardial infarction (MI) induced HF, as well as samples from ischaemic human HF patients (classified as reduced ejection fraction, details in Supplementary Table 1). In contrast to LVH, zDHHC5 expression was unchanged or modestly reduced in post-MI samples compared to control. Additionally, despite RNA sequencing data suggesting zDHHC5 expression levels are reduced in HFpEF but unchanged in HFrEF (Figure 1; Hahn *et al*., 2021) we found that protein expression of zDHHC5 was also significantly reduced in ischaemic HF samples compared to organ donors (Figure 5). We also investigated the expression of thioesterase APT1, which depalmitoylates NCX1, but found there was no significant change in expression in these samples (Supplementary Figure 4; Gök *et al*., 2021).

**Figure 5:**
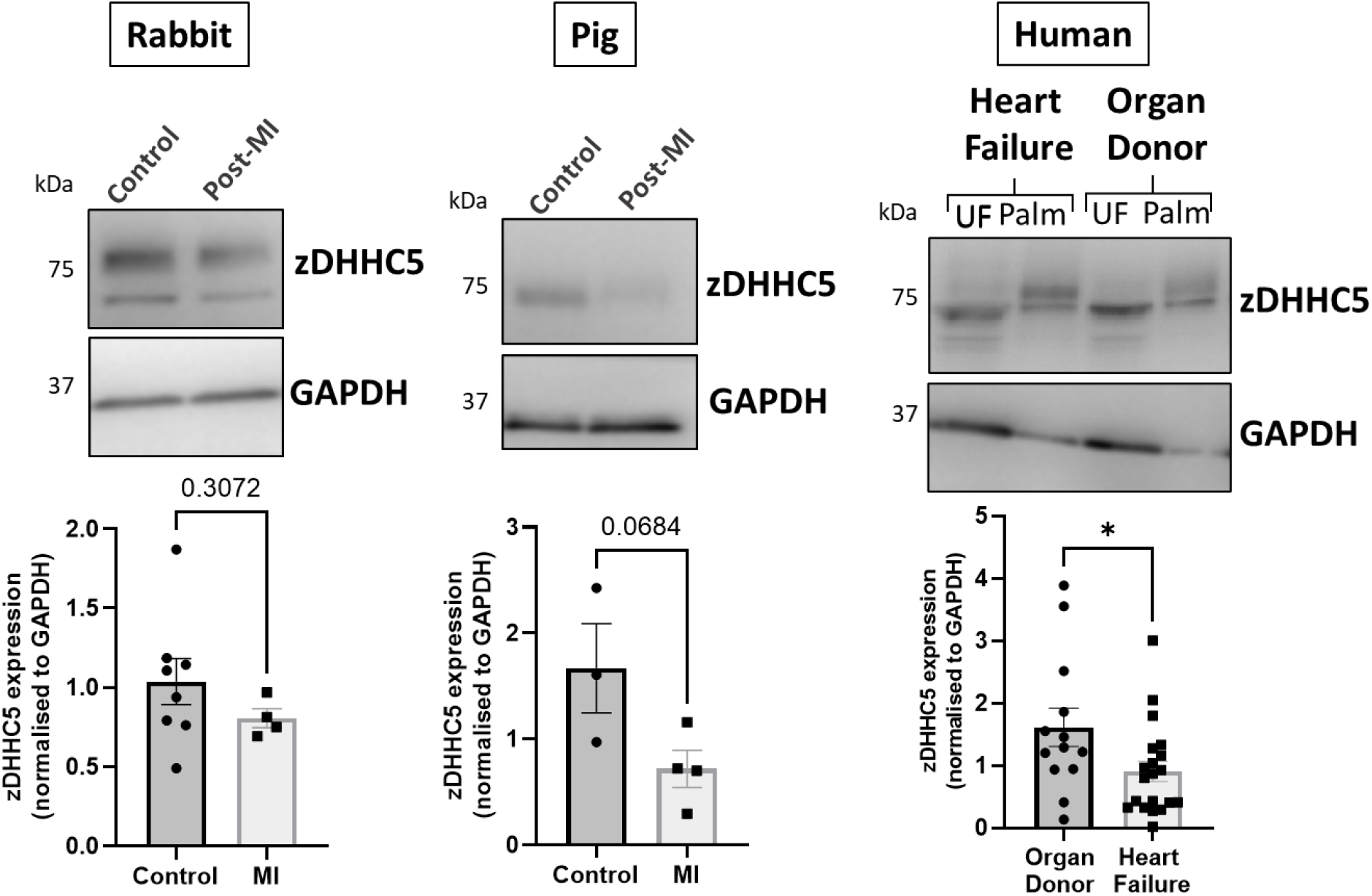
zDHHC5 expression in heart failure. In the rabbit model of MI-induced HF (8-weeks post-MI), zDHHC5 expression was unchanged whilst in the pig model of MI-induced HF with reperfusion (3-months post-MI) zDHHC5 expression was modestly reduced (p=0.0684). In samples from patients with ischaemic heart failure, zDHHC5 expression was significantly reduced compared to organ donor controls. zDHHC5 expression was normalised to loading control GAPDH. Statistical comparisons made by unpaired Student’s t-test. Data are mean ±S.E.M. *p<0.05.

Given both animal models and human HF samples were associated with reduced zDHHC5 expression, we investigated whether this correlated with a change in the palmitoylation of zDHHC5 substrates NCX1 and PLM. Whilst PLM palmitoylation remained unchanged in all cases, NCX1 palmitoylation was significantly reduced in animal models of HF but increased in human heart failure (Figure 6).

**Figure 6:**
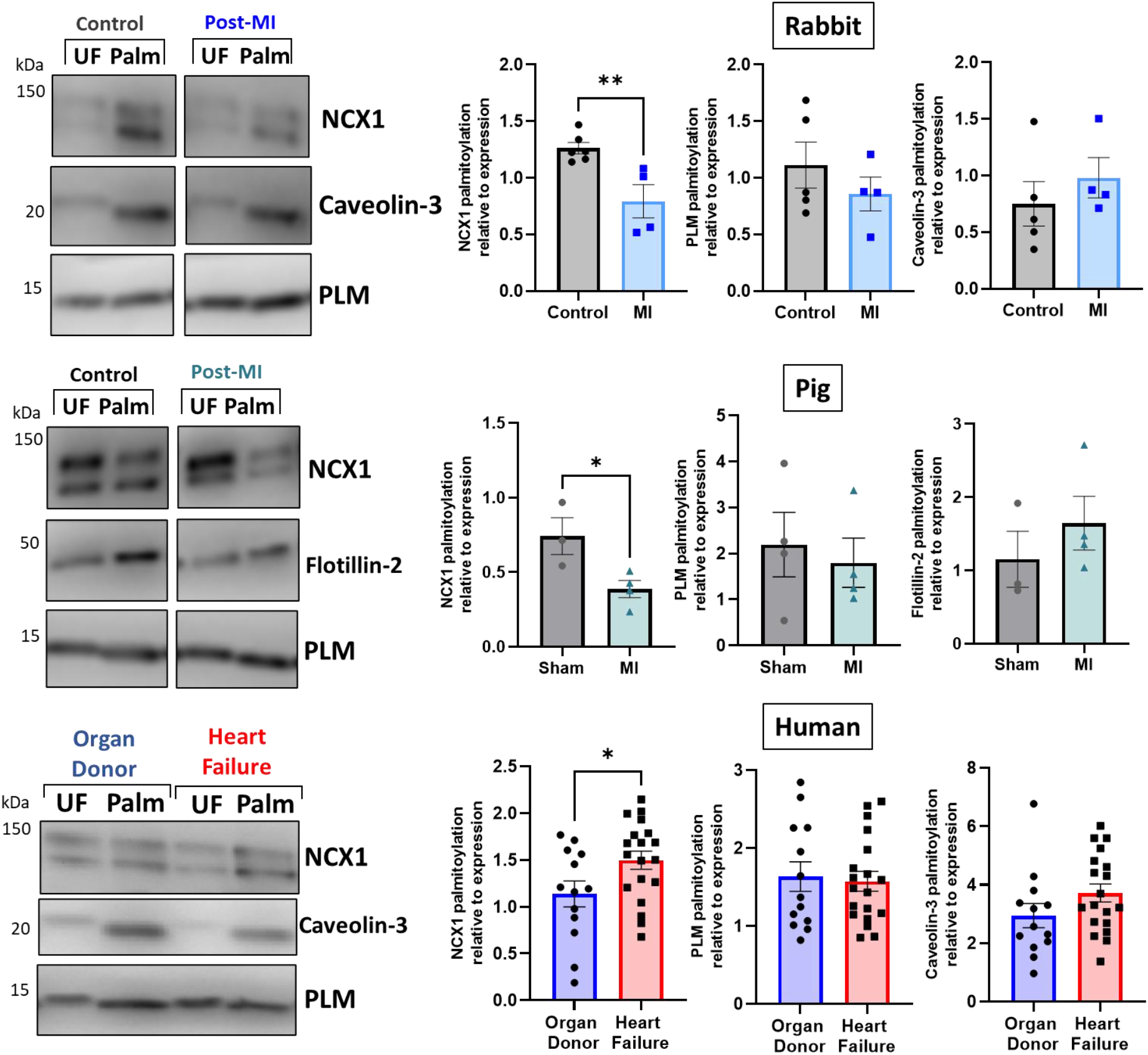
Palmitoylation of zDHHC5 substrates NCX1 and PLM in heart failure. Palmitoylation of NCX1 and PLM was investigated in tissue from a rabbit model (8-weeks post-MI) and a pig model (3-months post-MI) of experimental HF, as well as samples from human organ donor and ischaemic heart failure patients. Palmitoylation of NCX1 was significantly reduced in animal models of HF but increased human heart failure samples, whilst palmitoylation of PLM and assay control Caveolin-3/Flotillin-2 remained unchanged. Data are palmitoylated fraction normalised to total protein (unfractionated, UF). Statistical comparisons made by unpaired Student’s t-test. Data are mean ±S.E.M. *p<0.05, **p<0.01.

In the present study, zDHHC5 expression levels appear to correlate poorly with palmitoylation of its substrates, suggesting changes in zDHHC5 expression may not be sufficient to increase palmitoylation of its substrates. ZDHHC5 is the target of PTMs itself, including palmitoylation, which occurs on its active site cysteine during autopalmitoylation before transferring the palmitate to a substrate cysteine (Brigidi *et al*., 2015). Additionally, zDHHC5 is palmitoylated at additional sites in its C-terminal tail, which is key to mediating its response to β-adrenergic signalling and facilitating its interaction with the Na^+^/K^+^ ATPase, which regulates recruitment and palmitoylation of PLM (Chen *et al*., 2020; Plain *et al*., 2020; Yang *et al*., 2010),. As such, we investigated whether palmitoylation of zDHHC5 was changed in HF. Interestingly, zDHHC5 palmitoylation was altered in HF in a similar manner to that of NCX1, whereby zDHHC5 palmitoylation was significantly reduced in the pig model (Figure 7A), but modestly increased in human HF samples, although this change is not significant (Figure 7B).

**Figure 7:**
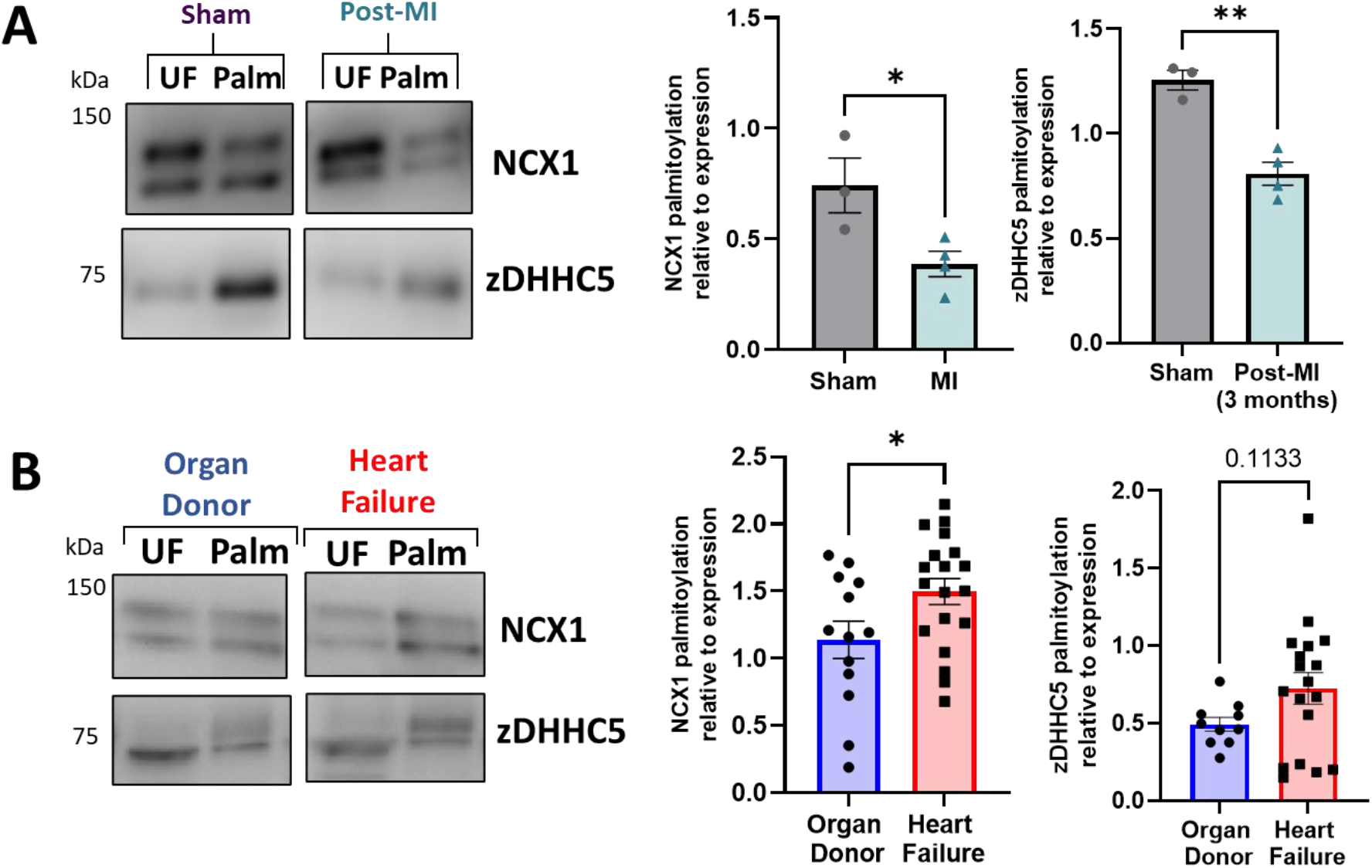
zDHHC5 palmitoylation is altered in heart failure in a similar manner to NCX1 palmitoylation. **A)** In a pig model of MI-induced heart failure, zDHHC5 palmitoylation is significantly reduced, whilst **B)** in human heart failure samples, zDHHC5 palmitoylation is modestly increased (p=0.1133). NCX1 data taken from Figure 6 for comparison. Statistical comparisons made by unpaired Student’s t-test. Data are mean ±S.E.M. *p<0.05, **p<0.01.

## Discussion

Palmitoylation has emerged over the last decade a crucial regulatory modification for every class of protein, including several involved in cardiac excitation-contraction coupling (Chien *et al*., 1996; Tulloch *et al*., 2011; Howie *et al*., 2013; Gök., 2020; Main *et al*., 2022). As many of these proteins are dysregulated in diseases such as HF, palmitoylation may represent a novel mechanism to manipulate their function for therapeutic benefit (Main & Fuller, 2021). Despite this, studies investigating changes in palmitoylation in a cardiac disease setting have been limited. As such, we focussed our investigation on the most well classified cardiac zDHHC-PAT, zDHHC5, which has been implicated in A/R injury and regulates the palmitoylation of important ion transporters and accessory proteins in the heart (Tulloch *et al*., 2011; Hilgemann *et al*., 2013; Lin *et al*., 2013; Howie *et al*., 2014; Chen *et al*., 2020; Gök *et al*., 2020; Plain *et al*., 2020). In the present study, we provide the first evidence that zDHHC5 expression and palmitoylation are altered in cardiac disease, although this does not directly correlate with a change in the palmitoylation of its substrates NCX1 and PLM.

Firstly, we observed that in a LVH model of cardiac hypertrophy and remodelling, zDHHC5 expression was upregulated as early as 3 days post-onset, and this was maintained until 8 weeks post-injury (Figure 2). Because there was such an acute upregulation of zDHHC5, we virally over-expressed zDHHC5 in cardiomyocytes to determine if there was a functional consequence for ventricular contractility, but this had no effect on any contractile parameters investigated (Figure 3). In both LVH samples and virally infected cardiomyocytes, despite increased zDHHC5 expression, there was no correlated increase in substrate palmitoylation, as palmitoylation of PLM remained unchanged, whilst NCX1 palmitoylation was significantly reduced in LVH samples (Figure 4). This suggests that zDHHC5 expression levels are not rate limiting for substrate palmitoylation in ventricular muscle. In other settings, availability of palmitoyl-CoA (Lin *et al*., 2013) and activity of acyl-CoA synthase enzymes (Gök *et al*., 2022), which may associate with zDHHC-PATs (Plain *et al*., 2020), have been established to be important determinants of substrate palmitoylation.

We investigated whether the changes in zDHHC5 expression were observed later in disease pathogenesis using models of ischaemic HF, including a rabbit MI model and an ischaemia/reperfusion pig model, as well as samples from ischaemic HF patients. In contrast to cardiac hypertrophy, zDHHC5 expression was either unchanged (rabbit), modestly reduced (pig) or significantly reduced (human) in a HF setting (Figure 5). However, similarly to the LVH model, this was poorly correlated with changes in substrate palmitoylation, whereby palmitoylation of NCX1 was significantly reduced in the animal models of HF, but was significantly increased in human HF samples (Figure 6). Throughout our investigation we observed changes in zDHHC5 expression that did not consistently match changes in palmitoylation its substrate NCX1. ZDHHC5 itself is under the control of several regulatory pathways, including palmitoylation of cysteines in its C-terminal tail which has important functional consequences for substrate recruitment and palmitoylation (Chen *et al*., 2020; Plain *et al*., 2020). In addition, its activity may be controlled by the availability of its acyl-CoA substrate, synthesised by ACSLs. Analysis of zDHHC5 palmitoylation revealed that, similar to NCX1, palmitoylation was significantly reduced in the pig model and modestly increased (although not significantly) in the human heart samples (Figure 7). This may suggest there are upstream regulatory pathways driving changes in substrate palmitoylation, including that of zDHHC5. ZDHHC-PATs have been frequently reported to palmitoylate each other, and a proximity biotinylation screen identified DHHC20 as an interactor and palmitoylating enzyme of zDHHC5. Palmitoylation of zDHHC5 C-terminal tail in response to adrenergic stimulation is required for its own stabilisation at the plasma membrane (Chen *et al*., 2020; Woodley & Collins, 2021). The zDHHC5 palmitoylation site lies in an amphipathic helix containing a binding site for the Na^+^/K^+^ ATPase and zDHHC5 accessory protein GOLGA7, which controls its membrane localisation (Plain *et al*., 2020; Woodley and Collins, 2019). Increased zDHHC5 palmitoylation as observed in human HF, or reduced palmitoylation as observed in the pig model, may suggest increased or decreased activity of zDHHC5 either through a change in palmitate loading into the active site or a change in zDHHC20-mediated palmitoylation. This is not uncommon, as although NCX1 expression is often increased in the setting of LV dysfunction in HF, this does not necessarily correlate with increased activity (Hobai & O’Rourke, 2000; Quinn *et al*., 2003).

Additionally, other PTMs of zDHHC5 will be important to consider in future investigations, as phosphorylation has been observed to inactivate zDHHC5 (Hao *et al*., 2020; Schianchi *et al*., 2020), while O-GlycNAcylation of DHHC5 enhanced PLM association and palmitoylation (Plain *et al*., 2020; Schianchi *et al*., 2020). In line with hypertrophy data, but in contrast to the ischaemic human HF results, Miles *et al*. report significantly increased levels of DHHC5 in human HF. However, the samples investigated represent a mixture of ischaemic and nonischaemic samples, limiting the comparison to the ischaemic samples used in this study (Miles *et al*., 2021). Nevertheless, given RNA-sequencing suggests levels will be significantly reduced in HFpEF and not HFrEF, further characterisation of zDHHC5 expression should be carried out in additional cohorts of patients (Hahn *et al*., 2021). Steady state protein palmitoylation is controlled by the balanced activities of palmitoylating and depalmitoylating enzymes. Several depalmitoylating enzymes show altered expression in both HFrEF and HFpEF, although perhaps to not the same extent as the dysregulated of zDHHC-PAT expression (Figure 1). In comparison to zDHHC-PATs, relatively little is known about the regulation of these enzymes, although APT1 was recently demonstrated to depalmitoylate NCX1 (Gök *et al*., 2021). However, we did not find a significant change in APT1 protein abundance in the setting of HF, so this likely does not explain the changes in substrate palmitoylation observed (Supplementary Figure 4).

Aside from information of zDHHC5 expression and palmitoylation, this study provides the first evidence that NCX1 palmitoylation is changed in cardiac disease in animal models and humans. Increased NCX1 palmitoylation, as observed in human HF, would enhance inactivation and reduce Ca^2+^ efflux which, whilst improving systolic function, this would contribute to diastolic impairment, of which NCX1 is a key mediator (Kass *et al*., 2004). Interestingly, the pattern of NCX1 palmitoylation associated with HF that we report here was recapitulated for cardiac myosin binding protein-C, again suggesting that upstream factors such as a fatty acid and fatty acyl-CoA availability may be regulating the changes of substrate palmitoylation (Main *et al*., 2022).

A significant finding, but also a limitation of this study, is the lack of consistency in the changes in protein palmitoylation between different animal models and human HF. This makes it challenging to draw definitive conclusions regarding the importance of palmitoylation in cardiac disease. The increased palmitoylation in the human setting may be a result of more developed decompensation compared to the animal models, or as a result of pharmacological intervention. It may also reflect a failure of the animal model to accurately reflect human pathology. Indeed, this is a major limitation of current animal models for HF research, because they rarely include study of novel therapeutic interventions in combination with current optimal clinical care, making translational potential challenging. Interestingly, whilst NCX1 palmitoylation was frequently altered, PLM palmitoylation did not change in any setting (Plain *et al*., 2020). However, this observation may be as a result of solely characterising palmitoylation using Acyl-RAC. Whilst this method provides a robust mechanism to detect protein palmitoylation, changes in substrate palmitoylation in singly palmitoylated proteins are most likely to be observed using this capture method, as opposed to substrates with multiple palmitoylation sites, where the substrate will still be captured even if only one site remains palmitoylated. Indeed, this may be why NCX1, with one palmitoylation site, is frequently observed to be changed whilst PLM containing two palmitoylation sites did not vary in any disease state (Howie *et al*., 2018; Reilly *et al*., 2015; Tulloch *et al*., 2011). Experimental approaches that measure palmitoylation site occupancy of multiply palmitoylated proteins may provide further insight into the contribution of aberrant protein palmitoylation to the pathogenesis of heart failure.

## Methods

### Human organ donor and ischaemic heart failure samples

Samples from organ donors (nonfailing) and human heart failure patients were obtained from the Gill Cardiovascular Biorepository at the University of Kentucky, of which patients and families of organ donors provided written consent. All procedures were approved by the local IRB and details of the collection procedures have been published previously (Blair *et al*., 2016). The study conformed with the principles in the Declaration of Helsinki. Samples in this study were taken from the ventricular endocardium of each heart with details in Supplementary Table 1.

### Mouse model of cardiac hypertrophy induced by pressure overload

Mouse samples were kindly provided by Professor Michael Shattock. Ventricular hypertrophy was induced by pressure overload via aortic constriction in 6-week old C57BL/6J mice, as has been previously described (Boguslavskyi *et al*., 2014).

### Porcine model of heart failure

Porcine samples were kindly provided by Dr. Roger Hajjar (New York, USA). Pigs were subjected to left anterior descending artery balloon occlusion to induce a myocardial infarction, as has been described previously (Tilemann *et al*., 2013).

### Ethical statement

Animals were handled in accordance with the UK Animals (Scientific Procedures) Act of 1986. All procedures were approved by the UK Home Office (PP7088535) and Glasgow University Ethics Review Committee. The animal research reported here adheres to the ARRIVE and Guide for the Care and Use of Laboratory Animals guidelines.

### Rabbit model of MI and heart failure

Adult male New Zealand White rabbits (12 weeks old, ∼3-4 kg) were given premedication with 0.4 mL/kg intramuscular Hypnorm (fentanyl citrate, 0.315 mg/mL: fluanisone 10 mg/mL, Janssen Pharmaceuticals). Anaesthesia was induced with 0.25–0.5 mg/kg midazolam (Hypnovel, Roche) via a cannula in the marginal ear vein. Rabbits were intubated and ventilated using a Harvard small animal ventilator with a 1:1 mixture of nitrous oxide and oxygen containing 1% halothane at a tidal volume of 50 mL and a frequency of 40 min^-1^. Preoperative antibiotic prophylaxis was given with 1ml Amfipen (ampicillin 100 mg/mL, Mycofarm UK Ltd) intramuscularly. A left thoracotomy was performed through the 4^th^ intercostal space. Quinidine hydrochloride (10 mg/kg; Sigma Pharmaceuticals), a class IA antiarrhythmic (potassium channel blocker) was administered intravenously prior to coronary artery ligation to reduce the incidence of ventricular fibrillation. The marginal branch of the left circumflex coronary artery, which supplies most of the LV free wall, was ligated halfway between the atrio-ventricular groove and the cardiac apex to produce an ischaemic area of 30–40% of the LV (Nisbet *et al*., 2016). Animals were maintained for 8 weeks to induce ischaemic cardiomyopathy phenotype which was confirmed by echocardiography measurements.

### Rabbit cardiomyocyte isolation

Adult rabbit ventricular myocytes (ARVM) were isolated from male New Zealand white rabbits as previously described. Isolation of adult rabbit ventricular cardiomyocytes (ARVM) was completed as described previously (Kettlewell *et al*., 2013). Briefly, New Zealand White male rabbits (12 weeks old, ∼3-4kg) were euthanised with a terminal dose of sodium pentobarbital (100mg/kg) with heparin (500IU), following which the heart was removed and retrogradely perfused on a Langendorff system. Enzymatic digestion of the tissue using a protease and collagenase solution occurred for ∼15 minutes before heart was removed from the system and cut into sections (left atria, right atria, right ventricle, left ventricle and septal regions) and each was finely dissected in Krafte-Brühe solution. The mixture was then triturated and agitated for 10 minutes before filtering and the cell suspension was centrifuged manually for a minute before the pellet was re-suspended in fresh KB. For experiments, cells were stepped up to physiological calcium in modified Krebs-Henseleit solution, initially containing 100µM of CaCl_2_ and left to settle for before the process was repeated using 200µM, 500µM, 1mM and 1.8mM concentrations of CaCl_2_. Cells were then snap frozen and kept at -80°C or used for functional experiments.

### Viral infection

Adenoviruses expressing zDHHC5 and zDHHS5 were produced in house using the AdEasy system (Agilent). Rabbit ARVM were infected for 18-24 hours by adding virus directly to the culture medium.

### Acyl-resin assisted capture

Acyl-resin assisted capture was used to purify palmitoylated proteins in a sample and adapted from a method published previously(Forrester *et al*., 2011). Cultured and pelleted cells were lysed in blocking buffer containing 1% methyl methanethiosulfonate (MMTS) to methylate free cysteines. Proteins were then precipitated using acetone and the resulting pellet subsequently washed using 70% acetone to remove excess MMTS. The pellets were then re-suspended in binding buffer before a portion of the sample was taken (unfractionated sample, total protein). To the remaining solution, 250mM NH_2_OH (hydroxylamine (HA), pH 7.5) was added to hydrolyse thioester bonds, with the same concentration of sodium chloride (NaCl, pH 7.5) added in its place for negative control samples. The proteins with free cysteines were the purified using thiopropyl sepharose beads. Palmitoylation of substrates was determined by relative quantity in HA samples compared to unfractionated.

### Immunoblotting

Standard western blotting was carried out using 6-20% gradient gels. The primary antibodies used were as follows: DHHC5 (1:1000, Sigma, HPA014670), NCX1 (1:1000, Swant, R3F1), PLM (FXYD1, 1:1000, Abcam, ab76597), Flotillin-2 (1:1000, BD Biosciences, 610383), Caveolin-3 (1:4000, BD Biosciences, 610420), HA-tag (1:5000, Roche, 11867423001). The secondary antibodies used were as follows: Rabbit anti-mouse HRP (1:2000, Jackson ImmunoResearch 111-035-144), Goat anti-rabbit HRP (1:2000, Jackson ImmunoResearch 315-035-003), Goat anti-rat HRP (1:2000, Jackson ImmunoResearch, 313-035-003), Donkey anti-guinea pig (1:2000, Jackson ImmunoResearch, 106-035-003).

### Confocal microscopy

Cardiomyocytes on 16mm glass coverslips were fixed with 4% paraformaldehyde (PFA) for 15 minutes at room temperature. Cells were permeabilised with 0.1% Triton-X100 in PBS for 10 minutes at room temperature before blocking with 3% BSA in PBS for at least 1 hour. Coverslips were incubated with HA-tag primary antibody (1:200, Roche, 11867423001) in 0.1% BSA in PBS for 1 hour before subsequent incubation with antirat Alexa Fluor 546 secondary antibody (1:400, Thermofisher, A-11081) for an additional hour. Coverslips were then mounted onto glass slides using Dako Fluorescence Mounting Medium with 1µl/ml 4’,6-diamidino-2-phenylindole (DAPI). Cells were then visualised using a Zeiss LSM 510 META Confocal Microscope with a 40x objective.

### Contractility measurements

CellOPTIQ® is an *in vitro* system designed by Clyde Biosciences which allows measurements of contractility, voltage and calcium to be carried out in individual adult cardiomyocytes (Clyde Biosciences Ltd; Glasgow, UK). Following culture and infection of cardiomyocytes for 18-24 hours, cells were transferred to a modified Krebs-Henseleit solution containing 1.8mM CaCl2 and incubated at 37°C using a heated stage. Cells were paced using electrodes with 40V pulses of 0.2ms duration at a frequency of 2Hz. Five second recordings at 100fps were then taken of individual cells using a 60x objective. Contractility recordings were analysed using an ImageJ Macro prepared by Dr Francis Burton which analyses the changes in sarcomere length as a measurement of cardiomyocyte contractility by determining the spatial frequency of the intensity profile of sarcomere bands over time. Each recording produced roughly 10 contractile peaks which were averaged to give one trace per cell which was analysed for contractility parameters of amplitude, time to up and down 90% of peak, contractile duration at 50% and 90% of peak and overall time to peak (Rocchetti *et al*., 2014).

### Statistics

Statistical analysis was completed using GraphPad Prism (Version 7; California, USA) and was performed on groups with 3 biological replicates or more. For comparisons in data sets with more than two groups, a one-way analysis of variance (ANOVA) with a Sidak’s or Dunnett’s post-hoc test was used, with comparisons detailed in the figure legend. For comparisons of two groups, a paired or unpaired Student’s t-test was used. All samples were tested for the presence of significant outliers (Rout’s test). A probability of p<0.05 was considered to be statistically significant.

## Acknowledgements

We acknowledge the patients and families of organ donors who donated cardiac samples. Funding: American Heart Association TP135689, NIH1L48785, University of Kentucky Myocardial Recovery Alliance.

## Availability of data and material

Further information and requests for resources and reagents should be directed to and will be fulfilled by the corresponding author, William Fuller

## Authors’ contributions

AM: Conceptualization, Investigation, Formal Analysis, Writing – original draft

WF: Conceptualization, Project Administration, Funding Acquisition, Supervision, Writing – review and editing

AB, JH, CK, AR: Resources and methodology

RH, FB, GLS, GSB, KSC, MJS: Resources

