## Supplementary Data for "zDHHC5 expression is increased in cardiac hypertrophy and reduced in heart failure but this does not correlate with changes in substrate palmitoylation"

**Supplementary Figures:**

**Human heart failure patient and organ donor details:**

| Record ID | Case type | Sex | Primary diagnosis |
|---|---|---|---|
| 24713 | Organ Donor | Female | N/A |
| 2B487 | Organ Donor | Male | N/A |
| 31331 | Organ Donor | Female | N/A |
| 4B3FA | Organ Donor | Female | N/A |
| 4D931 | Organ Donor | Male | N/A |
| 5155D | Organ Donor | Male | N/A |
| 632FD | Organ Donor | Male | N/A |
| 8CB30 | Organ Donor | Female | N/A |
| B23E3 | Organ Donor | Male | N/A |
| BC90C | Organ Donor | Female | N/A |
| DOF54 | Organ Donor | Male | N/A |
| D61ZE | Organ Donor | Male | N/A |
| FC3CB | Organ Donor | Female | N/A |
| 046E | Heart Failure | Male | Ischaemic cardiomyopathy |
| 05FF7 | Heart Failure | Female | Ischaemic HFrEF |
| 14C39 | Heart Failure | Male | Ischaemic cardiomyopathy |
| 3F6DC | Heart Failure | Male | Ischaemic cardiomyopathy |
| 58545F | Heart Failure | Male | Ischaemic cardiomyopathy s/p MI |
| 6DB85 | Heart Failure | Male | HFrEF from Ischemic cardiomyopathy |
| 7CE52 | Heart Failure | Female | Ischaemic heart failure |
| 8296A | Heart Failure | Male | Ischaemic cardiomyopathy |
| 8E8D8 | Heart Failure | Male | Ischaemic cardiomyopathy |
| 97CDC | Heart Failure | Male | Ischaemic cardiomyopathy |
| 9D7E9 | Heart Failure | Male | Ischaemic cardiomyopathy |
| AF1FF | Heart Failure | Male | Chronic systolic HF |
| B8BE2 | Heart Failure | Male | Ischaemic heart failure |
| BO644 | Heart Failure | Female | Ischaemic cardiomyopathy |
| C3B57 | Heart Failure | Male | Ischaemic cardiomyopathy |
| CB8A5 | Heart Failure | Male | Ischaemic cardiomyopathy |
| DA820 | Heart Failure | Male | Ischaemic cardiomyopathy |
| EF5CB | Heart Failure | Male | Ischaemic heart failure |
| FE8E2 | Heart Failure | Male | Chronic systolic HF |

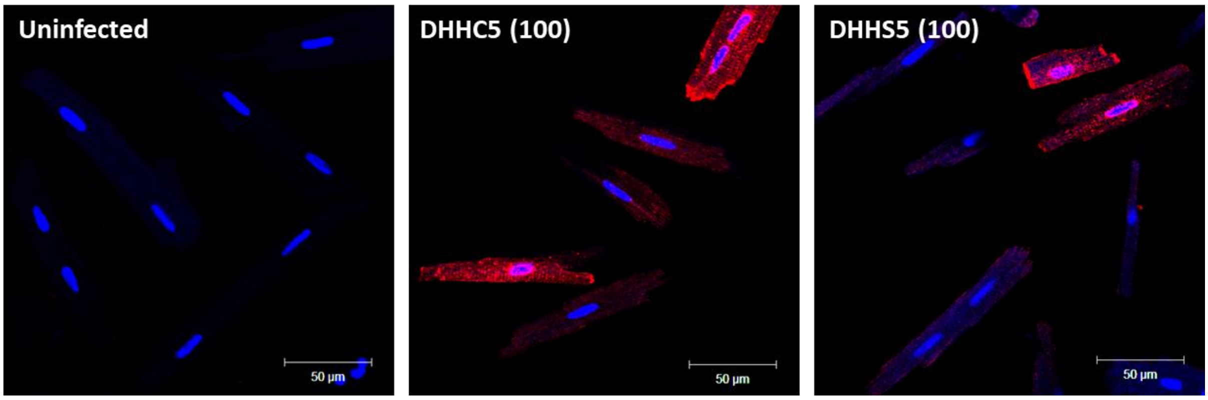


**Supplementary Figure 1: Virally infected HA-DHHC5 and HA-DHHS5 localises in intercalated discs, cellular membrane and perinuclear membrane in rabbit cardiomyocytes**. Rabbit ventricular cardiomyocytes were cultured for 18-24 hours in the presence of either HA-DHHC5 virus or a HA-DHHS5 dominant negative virus at a multiplicity of infection (MOI) of 100. Viral DHHC5 was detected using an anti-HA antibody, Alexa Fluor-546 secondary antibody and DAPI was used to stain the nuclei of the cells. Cells were visualised using a Zeiss LSM 510 META Confocal Microscope (40x objective) and 10 images per sample were captured. All cells appeared multinucleated and bright red fluorescence was observed at the intercalated discs, cell surface and perinuclear membrane in infected cells but not in uninfected controls. Representative images of the uninfected and viral samples are shown with contrast enhanced using ImageJ. Scale bar represents 50µm.

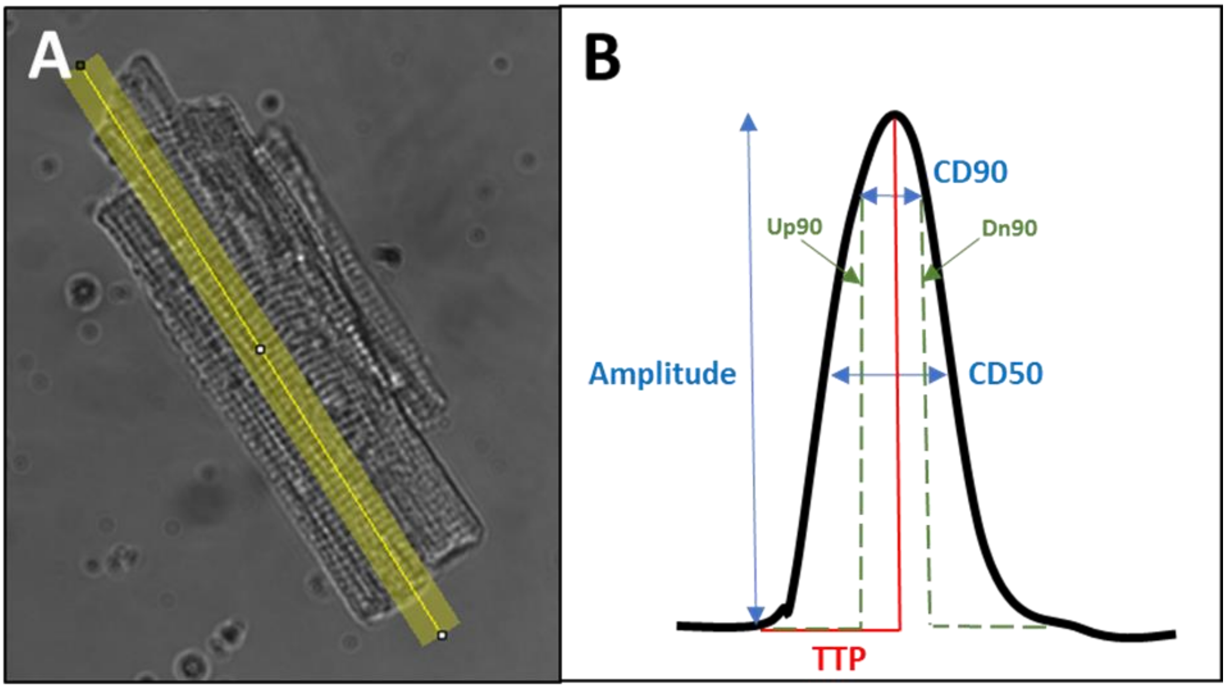


**Supplementary Figure 2: CellOPTIQ® measurement of cardiac contractility**. **A)** Cardiomyocyte contractility is recorded at 100fps and a section for analysis highlighted in ImageJ. **B)** An ImageJ Macro (Clydes Biosciences, Ltd) produces an average fractional shortening trace determined from changes sarcomere length over time. A number of parameters can be determined from this including amplitude of peak, time to peak (TTP), time at 90% of the upstroke (Up90) and downstroke (Dn90), and contraction duration at 90% (CD90) and 50% (CD50) of the peak.

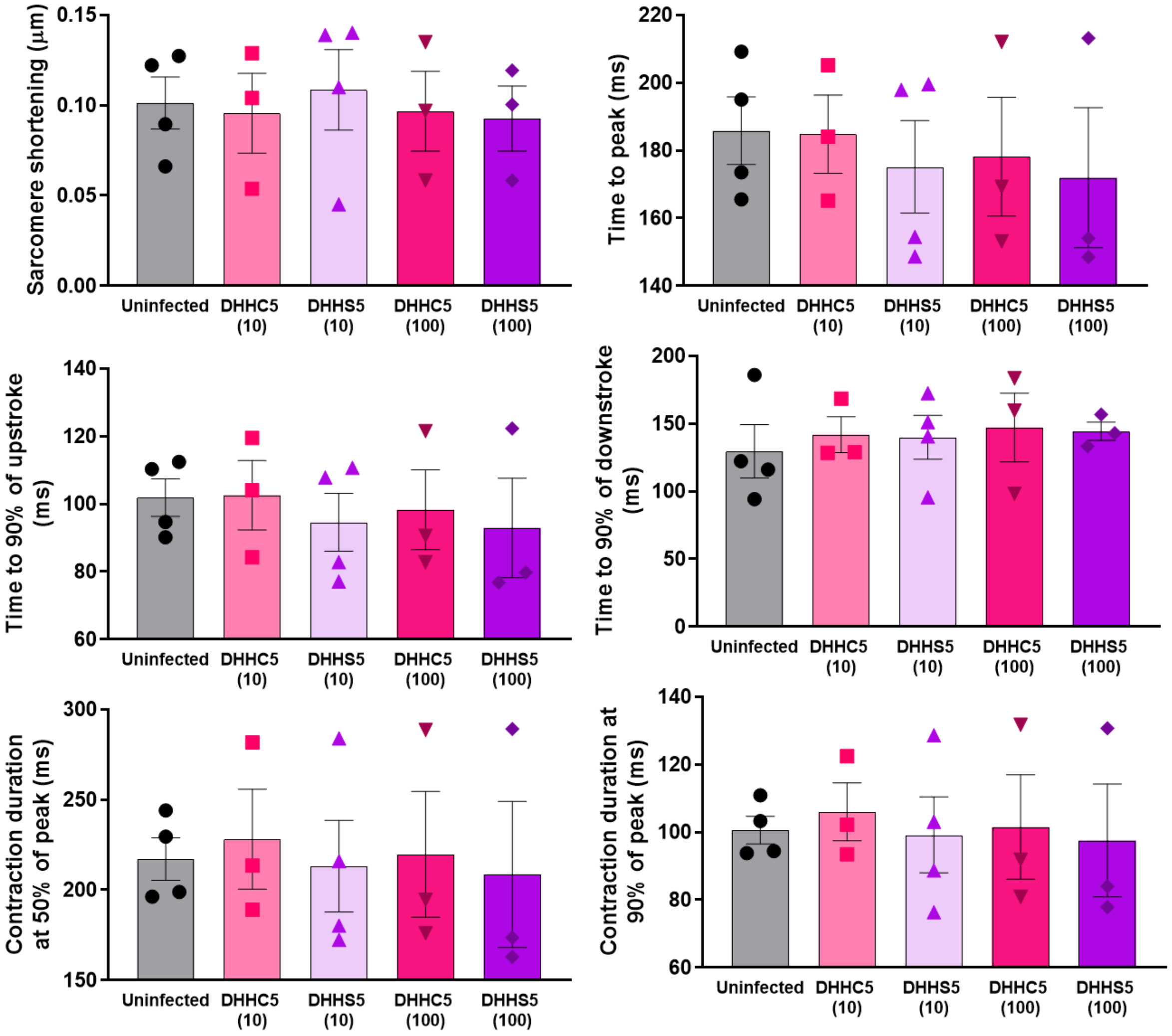


**Supplementary Figure 3: DHHC5 and DHHS5 overexpression do not alter parameters of rabbit ventricular cardiomyocyte contractility.** Rabbit septal cardiomyocyte cells were infected with DHHC5 or DHHS5 viruses at an MOI of 10 or 100 and cultured for 18-24 hours following which contractility recordings were taken using CellOPTIQ®, whereby cells were paced (2Hz frequency, 0.2ms duration, 40V) and 5 second recordings at 100fps were recorded. Recordings were analysed using an ImageJ macro and the contractility parameters of amplitude (sarcomere shortening), time to peak, time to 90% of peak upstroke, time to 90% of peak downstroke, contraction duration at 50% of peak and contraction duration at 90% of peak were compared across the groups. Viral infection with either DHHC5 or DHHS5 at MOI 10 or 100 had no significant effect on altering the parameters of contractility investigated. Data are mean ±S.E.M analysed using a one-way ANOVA with a Sidak's post-hoc test. N=3 biological replicates for S10, C100, S100 and n=4 biological replicates for uninfected and C100, with n=20-51 cells per replicate.

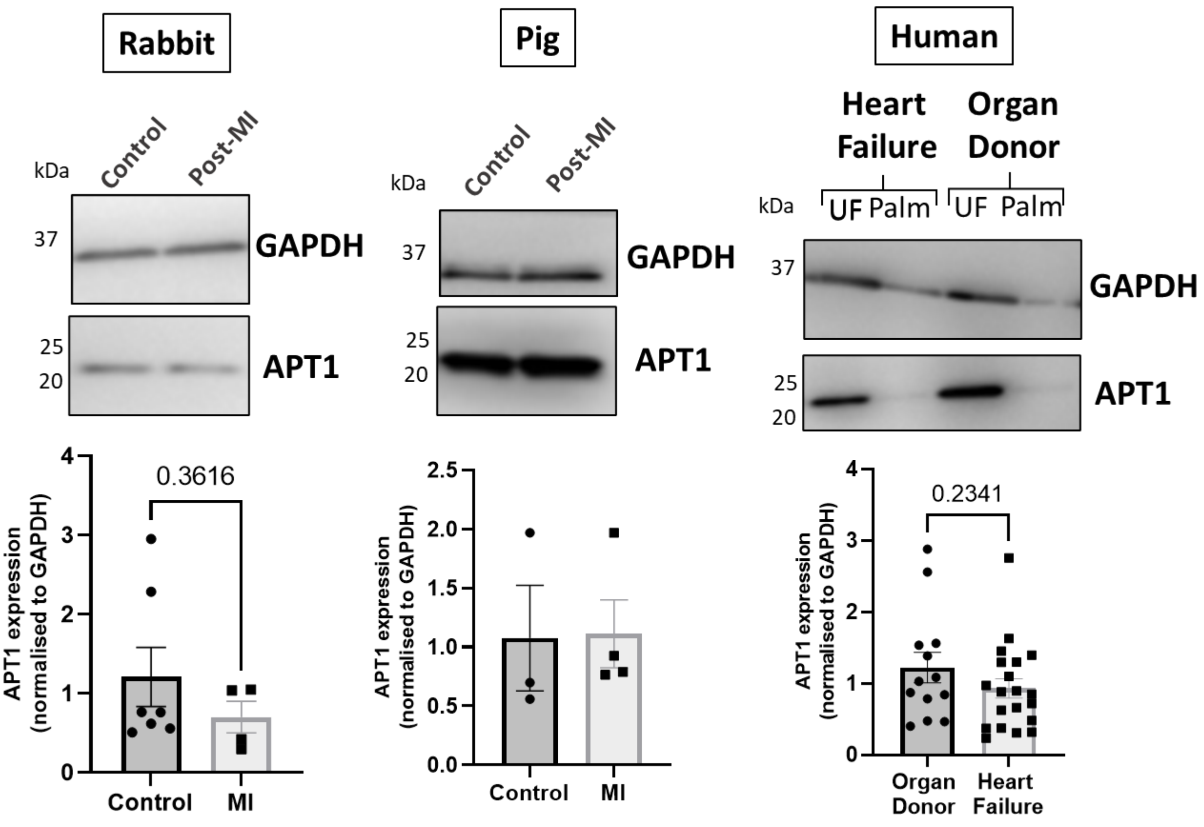


**Supplementary Figure 4: APT1 expression in heart failure.** In the rabbit model of MI-induced HF (8-weeks post-MI), a pig model of MI-induced HF with reperfusion (3-months post-MI) and samples from patients with ischaemic heart failure, APT1 expression was not significantly different to controls or organ donors. APT1 expression was normalised to loading control GAPDH. Statistical comparisons made by unpaired Student's t-test. Data are mean ±S.E.M**.**
